# Emergence and expansion of highly infectious spike:D614G mutant SARS-CoV-2 in central India

**DOI:** 10.1101/2020.09.15.297846

**Authors:** Shashi Sharma, Paban Kumar Dash, Sushil K Sharma, Ambuj Srivastava, Jyoti S Kumar, B.S. Karothia, K T Chelvam, Sandip Singh, Abhay Gupta, Ram Govind Yadav, Ruchi Yadav, TS Greshma, Pramod Kushwah, Ravi Bhushan, D.P. Nagar, Manvendra Nandan, Subodh Kumar, Duraipandian Thavaselvam, Devendra Kumar Dubey

**Author notes:** Corresponding author Dr Paban Kumar Dash, Dr Subodh Kumar, Email-, Defence R & D Establishment, Jhansi Road, Gwalior – 474002, India.

## Abstract

COVID 19 has emerged as global pandemic with largest damage to the economy and human psyche. The genomic signature deciphered during the ongoing pandemic period is valuable to understand the virus evolutionary patterns and spread across the globe. Increased availability of genome information of circulating strain in our country will enable to generate selective details in virulent and non virulent markers to prophylaxis and therapeutic interventions. The first case of SARS CoV-2 was detected in Chambal region of Madhya Pradesh state in mid of March 2020 followed by multiple introduction events and expansion of COVID-19 cases within 3 months in this region. We analyzed around 5000 COVID -19 suspected samples referred to Defence Research and Development Establishment, Gwalior, Madhya Pradesh. A total of 136 cases were found positive over a span of three months period this includes virus introduction to region and further spread. Whole genome sequences employing Oxford nanopore technology were deciphered for 26 SARS-CoV-2 circulating in 10 different districts in Madhya Pradesh State of India. The region witnessed index cases with multiple travel history responsible for introduction of COVID-19 followed by remarkable expansion of virus. The genome wide substitutions including in important viral proteins were observed. The detailed phylogenetic analysis revealed the circulating SARS-CoV-2 clustered in multiple clades A2a, A4 and B. The cluster wise segregation was observed suggesting multiple introduction links and evolution of virus in the region. This is the first comprehensive details of whole genome sequence analysis from central India region, which will add genome wide knowledge towards diagnostic and therapeutic interventions.

## Introduction

Emergence of novel virus is considered as a major challenge to humanity. SARS CoV-2, the etiology of “coronavirus disease 2019” (COVID-19) has emerged in Wuhan, China, in December 2019 (CSG, 2020). COVID-19 was declared as a public health emergency of international concern (PHEIC) by WHO in 30 January 2020. Subsequently, the incidence of disease and mortality increased across the globe. Finally, WHO declared COVID-19 as a pandemic on 11^th^ March, 2020 based on the speed and scale of transmission across the globe (Andersen et al., 2020). The disease has affected more than 17 million persons as on 30^th^ July 2020 and still rising rapidly. The common signs of infection include cough, fever, sore throat, respiratory symptoms inclusive of shortness/difficulties in breathing. More severe symptoms can include pneumonia, severe acute respiratory syndrome, kidney failure and even death with coalescence of factors (Zhu et al., 2020, Youg et al., 2020). Many COVID-19 cases have been reported to be asymptomatic and they serve as carrier of SARS-CoV-2 (Xu et al., 2020; He et al., 2020).

SARS-CoV-2, the etiological agent of COVID-19 is a novel virus and a member of subgenus-Sarbecovirus, genus-Betacoronavirus of family-*Coronaviridae*. SARS-CoV and MERS CoV, etiology of SARS and MERS are the closest relative within this group. Whole Genome Sequences (WGS) of SARS-CoV-2 suggest bat-CoV to be its closest progenitor with 96% homology. However, Receptor binding domain (RBD) of its Spike protein with an efficient binding to ACE-2, the receptor for SARS-CoV-2 in human cell seems to have been derived from Pangolin CoVs (Wan et al., 2020, Andersen et al., 2020).

The first three cases in India were reported from the state of Kerala in late January and early February, with a travel history of Wuhan, China. India took drastic steps to contain the further spread of the virus including imposition of travel restrictions to-and-from the affected countries. There were no new cases of COVID-19 for almost a month. All three cases subsequently tested negative making India free of the disease at that point of time (Press Information Bureau, PIB, India, 2020). However, while the global focus was on China and other eastern countries like South Korea and Japan; European countries, middle-east and the USA reported a surge in cases of COVID-19, pressing the WHO to declare it as a pandemic. Since March 2020 onwards, India also witnessed a surge of imported cases from countries other than China which has been further assisted with local transmission. In March, imposition of nationwide lockdown checked the epidemic curve. Despite these measurements, the exodus of individual from their place of work to native places led to rapid transmission of virus to remote corners of the country. India now stands dangerously at 3^rd^ position with overall 1.8 million reported infections with the cases accelerating rapidly.

The genome of SARS-CoV-2 comprises of a single stranded positive sense RNA of approximately 29.9kb long that codes for 14 ORFs and 27 proteins. Out of these, there are four structural proteins viz., spike surface glycoproteins (S), Envelope protein (E), Membrane protein (M) and Nucleocapsid protein (N); 15 non structural protein (nsp1-nsp10) and nsp12-nsp16 and 8 accessory proteins 3a, 3b, p6, 7a, 7b, 8b, 9b and ORF 14.

Genetic characterization of a virus is extremely important from both epidemiological and microbial forensics points of view. We at Defence Research & Development Establishment (DRDE) Gwalior being a nodal center for COVID-19 diagnosis engaged in screening of COVID-19 suspected cases from 10 different districts viz., Gwalior, Morena, Ashoknagar, Sheopur, Shivpuri, Guna, Datia, Bhind, Jabalpur, Bhopal. In this region the specimens were collected by medical authorities at different district hospitals from mid of March to May 2020. Around 5000 samples were referred with detailed patient history from the district health authorities to DRDE, Gwalior.

Next generation sequencing (NGS), an emerging technology aided understanding of evolution of SARS-CoV-2 genomes and its transmission patterns after it enters a new population (Chen and Li et al., 2020). This is proving to be an important step towards formulating strategies for management of this pandemic. In this study, we characterized the genome of representative viruses involved in index cases of the region along with expansion and fatal cases. Further the whole genome analysis was performed to annotate genome wide amino acid substitutions. The detailed molecular phylogentic analysis was carried out to classify viruses into different clusters and to understand their transmission linkages.

## Methods

### COVID-19 laboratory screening

Nasopharyngeal/Nasal/Oropharyngeal swabs in viral transport medium (VTM) received from acute phase patients with defined symptoms, asymptomatic cases with contact history with positive patients/ travel history were processed for laboratory confirmation of SARS-CoV-2 at Defence Research and Development Establishment, (DRDE) Gwalior, M.P., India. These samples were referred from 10 different districts health authorities of Madhya Pradesh, India during the period of March –May 2020. The laboratory diagnosis was accomplished through a Taqman qRT-PCR as recommended by WHO and ICMR.

### Isolation of viral RNA

The samples were processed in a BSL-3 facility (High Containment Facility) following biosafety precautions at DRDE, Gwalior. The viral RNA was extracted from the referred samples using QIAamp viral RNA mini kit (Qiagen, Germany) as per manufacturer’s instructions. Briefly, 560 µl of lysis buffer was added to 140 µl of clinical specimen and incubated for 10 min at room temperature, the sample was passed through silica column followed by washing in 500 µl wash buffer and finally the viral RNA was eluted in 50 µl of elution buffer in a nuclease free tube and stored at 4°C before addition to downstream molecular detection assay.

### Laboratory Investigation of COVID-19

Presence of SARS-CoV-2 was investigated by performing SARS-CoV-2 screening and confirmatory assays targeting Envelope, RdRP, ORF 1ab, N genes along with Human Rnase P as a housekeeping gene control to ensure RNA quality during sample collection as per WHO protocol. The Taqman qRT-PCR was performed with dual labeled hydrogenic probes labeled with FAM/VIC (E/Rnase P-Screening Assay), FAM/VIC (RdRP/ORF-Confirmatory Assay), N gene (Confirmatory assay). The fluorescence signals were recorded in ABI 7500 Dx Real time PCR (ABI, USA).

### Severe acute respiratory syndrome coronavirus 2 whole genome sequencing

The representative positive cases were selected on the basis of patients with travel history in the region (n=6), patients with different age group from similar contact history (n=10); introduction links from different areas to different families (n=4), COVID-19 positive death cases (n=2) and index cases from different districts (n=4). Based on above mentioned criteria, qRT-PCR positive cases (n=26) were selected for whole genome sequencing. Briefly cDNA was synthesized using Superscript IV reverse transcriptase (NEB, USA) along with 23 mer oligo dT linker, 50μM random hexamer and 10mM dNTPs mix. The reaction was incubated at 65°C for 5 min and snap chilled at ice, Further reverse transcription was performed at 42°C 50 min followed by 70°C 10min. The cDNA was stored at -80°C until further use.

### Amplification of Whole genome by Oxford Nanopore Platform

SARS CoV-2 whole genome sequencing was performed as per ARCTIC protocol by Josh Quick. Briefly cDNA from samples were used as template for multiplex PCR to amplify SARS-CoV-2 genome and the amplified products were purified. Purified cDNA amplicons from each sample were barcoded individually and pooled at equimolar concentration to prepare nanopore library. The quality and quantity of the pooled library was assessed using standard methods and sequenced on GridION-X5 nanopore sequencer. To generate tiled PCR amplicons from the SARS-CoV-2 viral cDNA, primers were designed using primal scheme. These primers were pooled into three different primer sets named as pool 1, 2 and 3.

Briefly multiplex PCR was set with initial denaturation at 98°C for 30sec followed by 25 cycles of denaturation at 98°C for 15 sec and annealing and extension at 65°C for 5 min. Amplification from respective primers sets were confirmed by agarose gel electrophoresis. Raw data generated was subjected to analysis using ARCTIC protocol to generate consensus sequences for SARS-CoV-2 genome.

### Nanopore library preparation and sequencing

Amplified products obtained from multiplex PCR of pool 1, 2 and 3 were pooled and purified by using Ampure-XP beads. 50ng from each samples were taken for library preparation using native barcoding kit and ligation kit from Oxford nanopore technology (ONT). Equimolar DNA from each sample was taken and end-repaired using NEBNext Ultra II End repair/da-tailing Module and cleaned with 0.4X AmPure xp Beads. Native barcode ligation was performed with NEBNext Ultra II Ligation Module using Native Barcoding kit. All barcode ligated samples were pooled together and purified using 0.4X of AmPure beads. Further sequencing adapter liagation was performed using NEB next quick ligation Module. Library mix was cleaned up using 0.4X AmPure beads and finally sequencing library was eluted in 15 μl of elution buffer and used for sequencing on SpotONflowcell in a 48hr sequencing protocol on GridION release 19.06.9. Nanopore raw reads (‘fast5” format) were basecalled (‘fastq5’ format) and multiplexed using Guppy v3.2.2.

### Data Analysis

Base-calling and demultiplexing of nanopore raw data was done using Guppy. The processed data was used for variant calling using Arctic pipeline and consensus generation using bam based method. The Arctic pipeline uses processed reads to align against the available SARS CoV-2 reference genome. Further, variant calling and consensus sequence generation was done using Nanopolish. The good quality variants were used for annotation using snpEff tool.

### Genome analysis of SARS-CoV-2

The nucleotide sequences of representative SARS-CoV-2 were retrieved from Global Initiative on Sharing All Influenza Data (GISAID) and NCBI GenBank, edited and analysed employing EditSeq and MegAlign modules of lasergene5 software package (DNASTAR Inc, USA). The complete genome of 26 SARS-CoV-2 deciphered in this study was comparatively analysed with prototype SARS CoV-2 isolated from Wuhan, China (GenBank Acc No MN908947). Multiple sequence alignment was carried out using MUSCLE alignment method in Bioedit software module. The amino acid substitutions of viruses sequenced in this study was compared to prototype SARS CoV-2 Wuhan strain.

Phylogenetic analysis based on SARS CoV-2 whole genomes were carried out with respect to globally diversified SARS CoV-2 (n=37) available at NCBI GenBank and GISAID data base from December 2019-May 2020 (Table 3). The phylogenetic tree was constructed employing Neighbor Joining method with 1,000 replicates of bootstrap analysis with general time-reversible model with gamma distributed rates of variations among sites using Mega 5.03 software.

**Table 1:** Ct values of samples sequenced in this study with respect to different gene targets detected by COVID-19 Taqman qRT-PCR.

| Sample ID | RdRP (Ct Value) | ORF 1 ab (Ct Value) | N2 (Ct Value) |
| --- | --- | --- | --- |
| 1 | 27 | 24.9 | 23.8 |
| 2 | 26.2 | 23.2 | 25.3 |
| 3 | 23.8 | 21.2 | 21.3 |
| 4 | 27.9 | 26.2 | 27.7 |
| 5 | 30.6 | 28.6 | 29.1 |
| 6 | 23.2 | 22.3 | 21.4 |
| 7 | 29 | 28 | 29 |
| 8 | 28 | 27.7 | 28.2 |
| 9 | 29 | 28 | 28 |
| 10 | 27.2 | 26.1 | 26.7 |
| 11 | 26.5 | 24.1 | 25.2 |
| 12 | 20.5 | 18.4 | 18 |
| 13 | 26.2 | 25 | 23 |
| 14 | 29.7 | 27 | 26.1 |
| 15 | 29.1 | 27.1 | 28 |
| 16 | 29.8 | 28.9 | 29.2 |
| 17 | 30.9 | 29.6 | 30 |
| 18 | 28.1 | 25.5 | 25.2 |
| 19 | 18.8 | 29.2 | 29 |
| 20 | 31.3 | 27.9 | 27.5 |
| 21 | 26.5 | 26.5 | 22 |
| 22 | 21.5 | 19.3 | 17.5 |
| 23 | 33.2 | 30.9 | 29.6 |
| 24 | 29 | 26.2 | 27.1 |
| 25 | 33.2 | 31.3 | 32.5 |
| 26 | 23.8 | 24.1 | 23.5 |

**Table 2:**
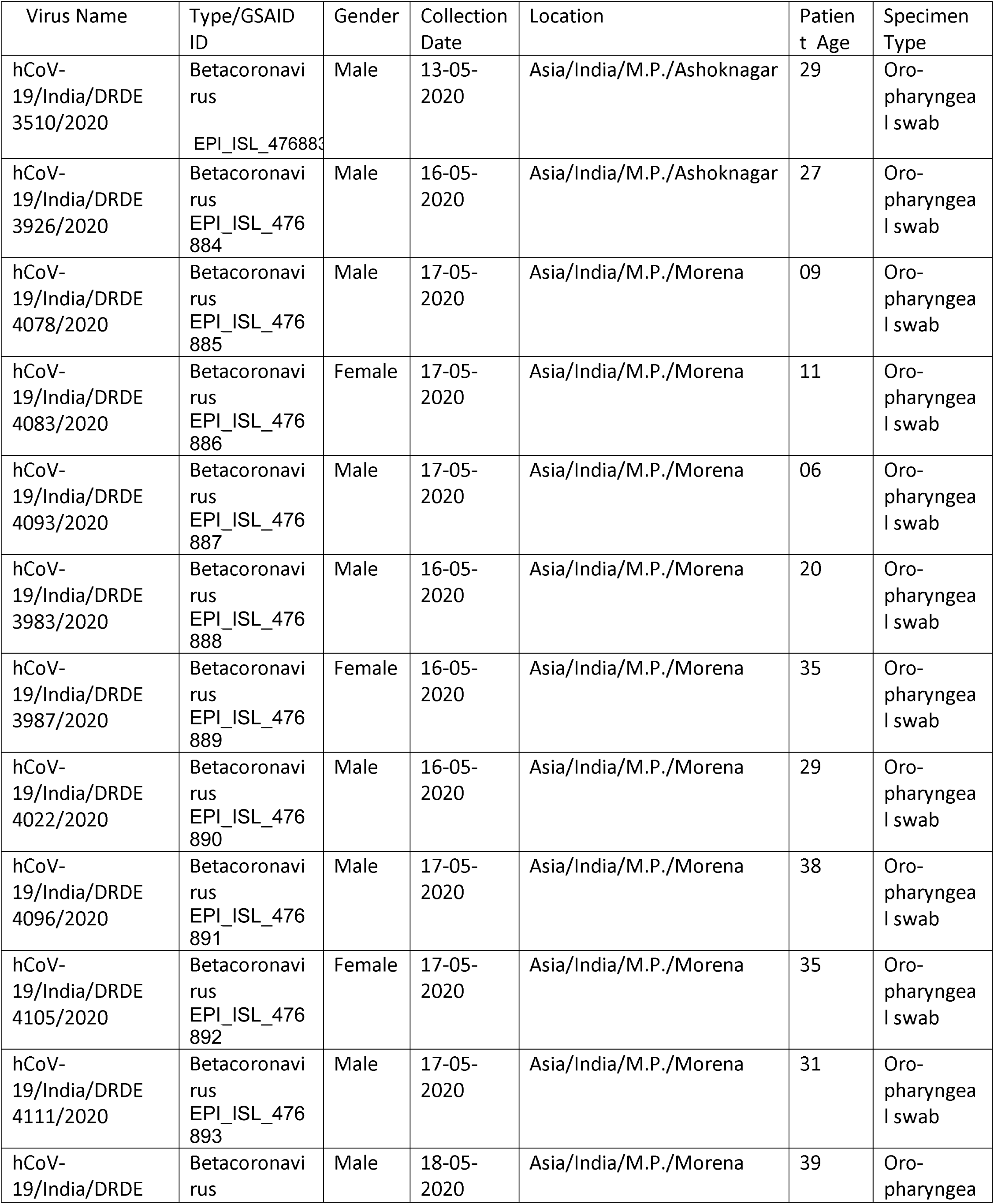
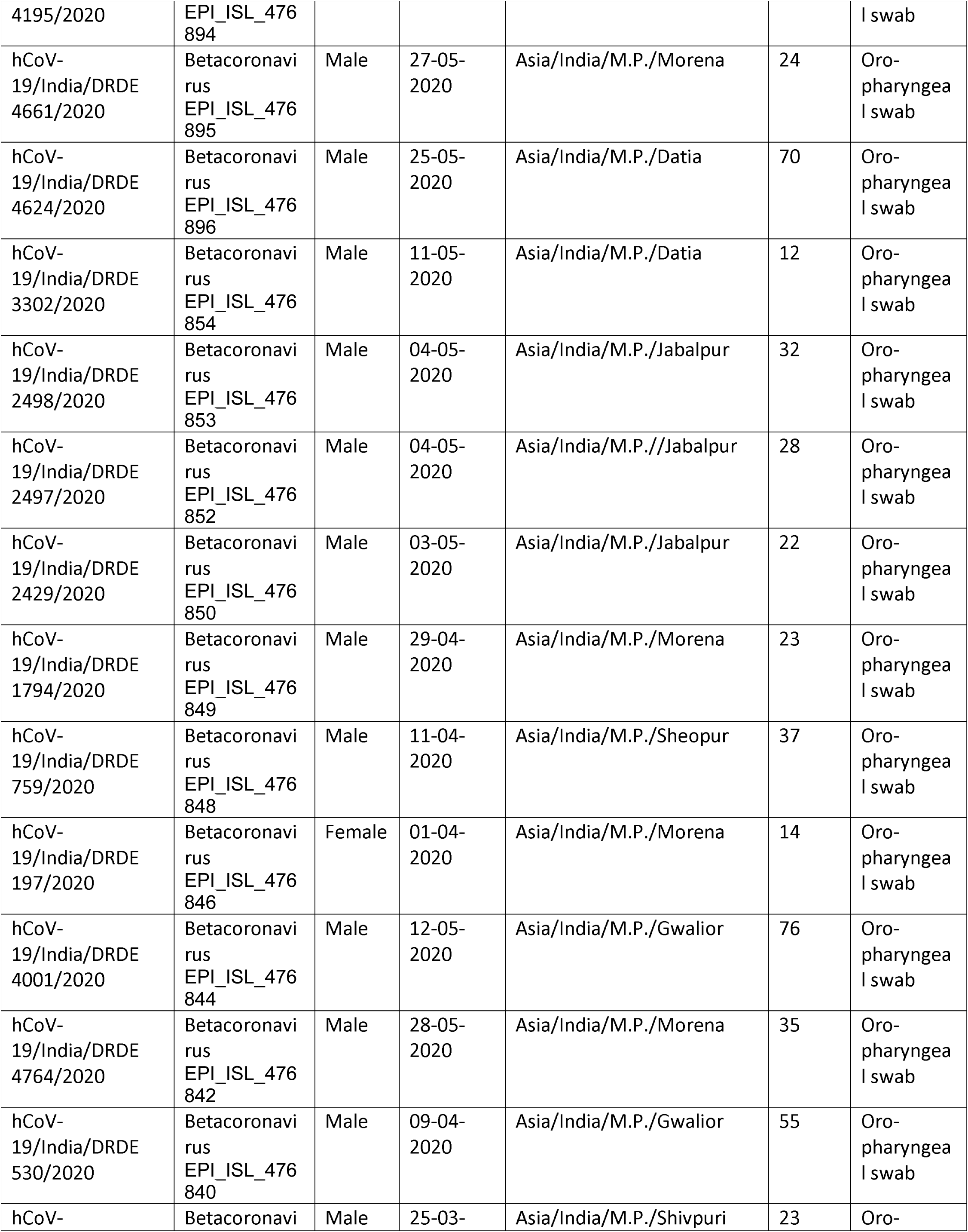
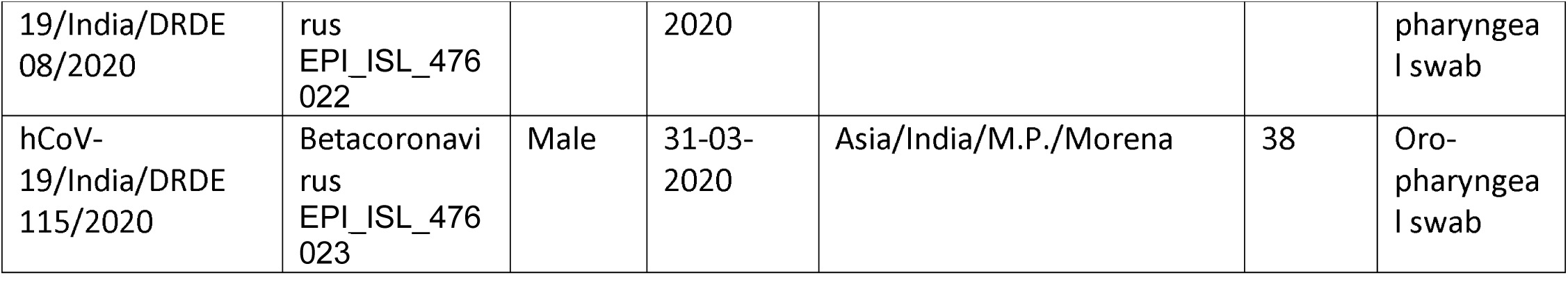
Indian SARS CoV-2 virus sequenced in this study.

**Table 3:**
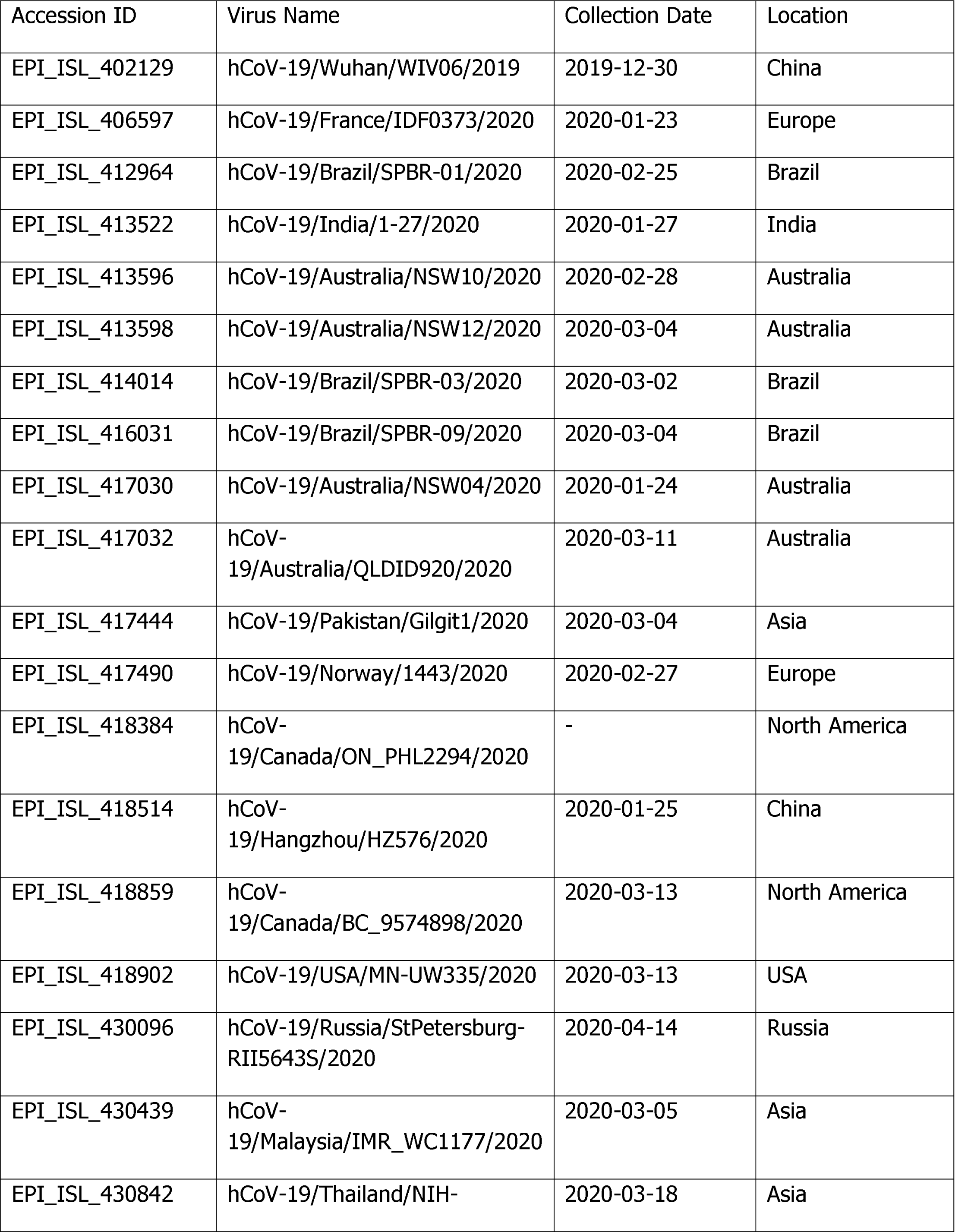
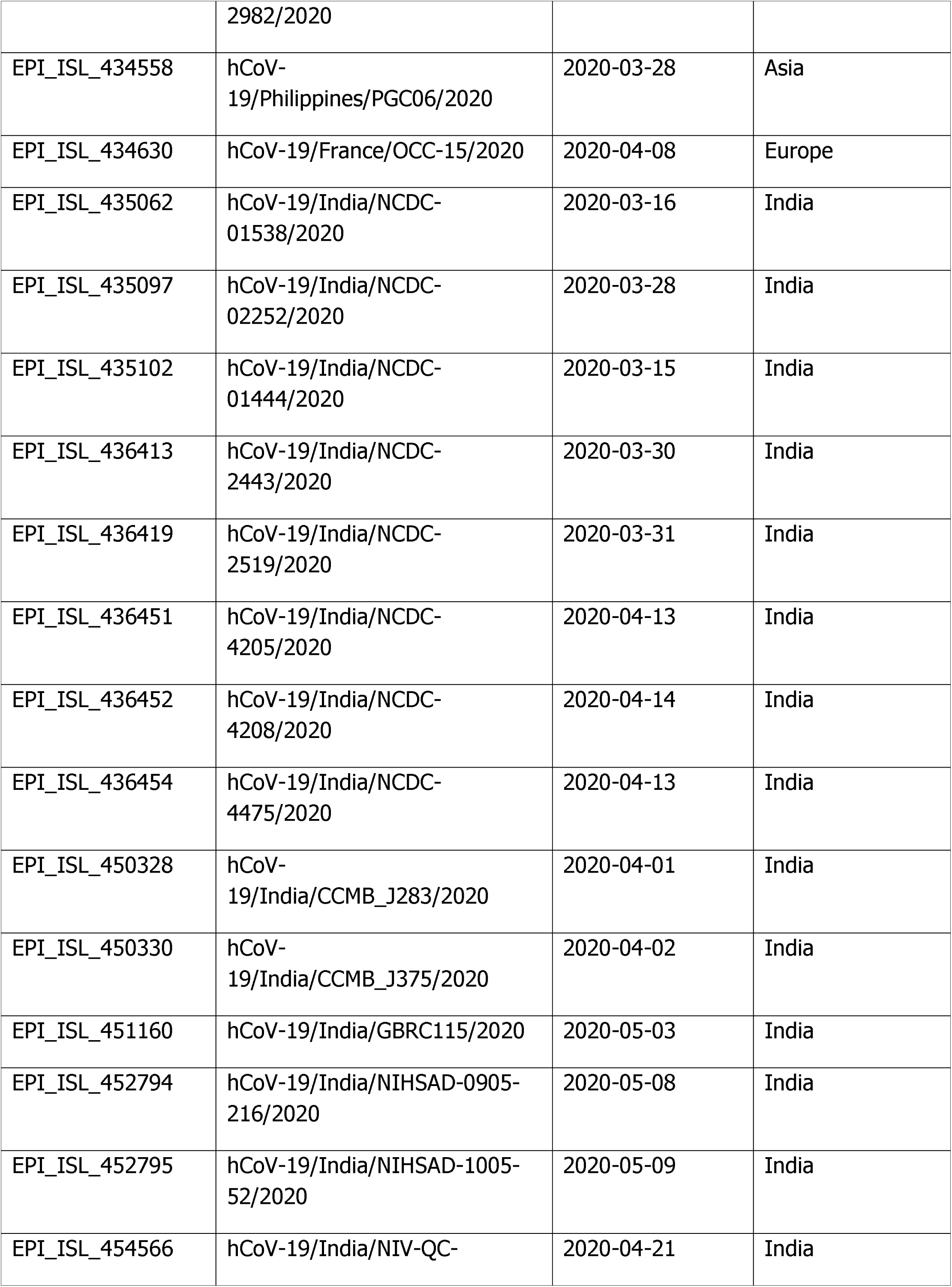
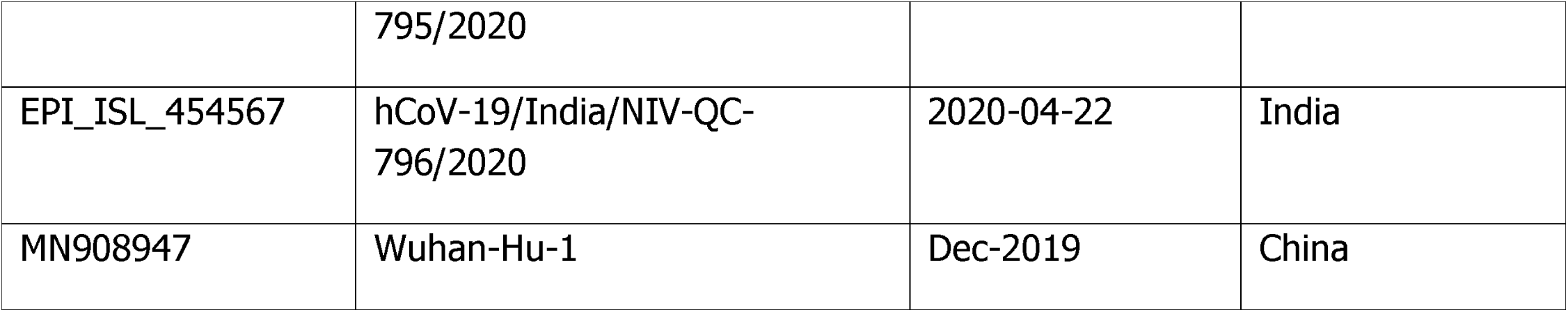
Details of Global SARS-CoV-2 virus sequences retrieved from GISAID web page.

## Results

### Clinical representation and laboratory diagnosis of COVID -19 cases

One hundred thirty six Cases referred to DRDE, Gwalior were found positive in period of three months (Last week of March-May 2020) diagnosed for COVID-19. All samples were investigated by WHO approved real time RT-PCR using SARS CoV-2 specific primers and probes for E, RP, RdRP and N genes. RNaseP as a housekeeping control was kept to ensure sample collection quality. All positive samples had shown clear positive amplification curve for presence of above mentioned genes (Fig. 1). The overall RT-PCR positivity was observed as 4%. The minimum age of COVID-19 infected cases were observed as 05 months whereas highest observed as 98 yrs. The maximum positive cases were found to belong to 21 to 30 year age group. Death in immunocompromised patients was reported within 24-48 hrs of hospitalization. The clinical history revealed the maximum symptomatic cases suffered from fever, sore throat, breathlessness. Maximum case positivity (68%) was reported from district Morena.

**Figure 1:**
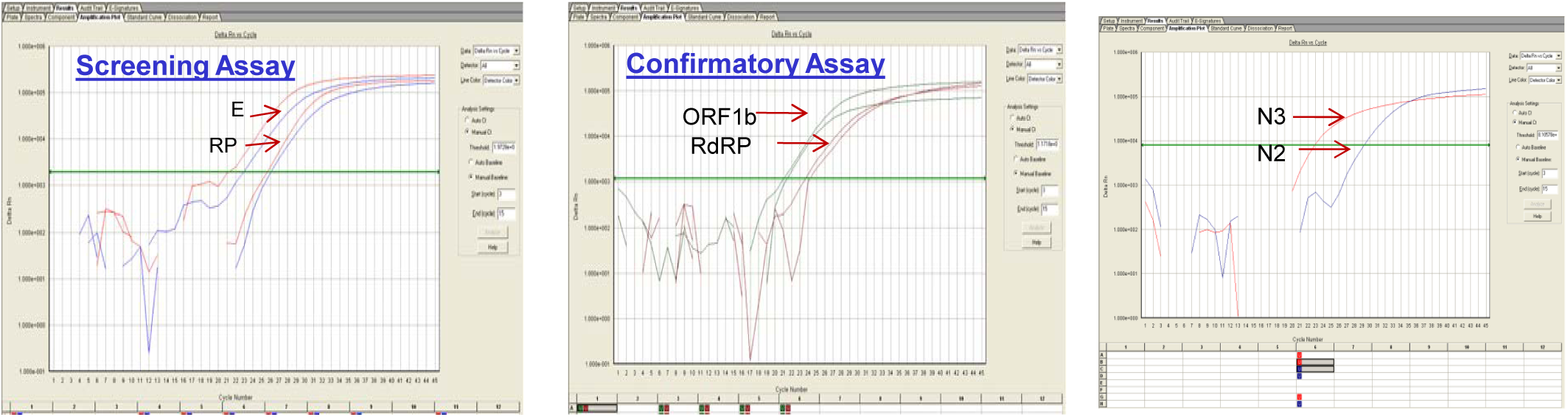
Amplification plot of positive samples showing amplification against different gene targets of SARS CoV-2 (E/RnaseP, Orf1b/RdRp, N genes)

### SARS-CoV-2 whole genome sequencing

Twenty six reference samples detailed in Table 3 along with GISAID accession numbers, patient age, location were sequenced in this study. The Ct values of all these 26 samples in respect to different genes were provided in Table 2. The viral RNA from above samples were converted to cDNA. The cDNA was processed for amplification through 3 pools of specific primers (Fig 2). The NGS led to generation of around 0.03 to 0.30 million nanopore reads. The length of RNA genomes was found to be 29.89 kb. The sequencing coverage for all samples was in range of 590X to 5000X showing high depth of sequence data. The processed reads from Guppy was used for variant calling using Arctic pipeline. The good quality filtered variants obtained was in the range of 5-13 across 24 samples. The predicted variants were further annotated using SnpEff TOOL. The reference genome of SARS CoV-2 was used for annotating good quality variants. The alignment files from Arctic pipeline were used for generating consensus sequence based on read depth of 10 supporting a position across genome. The bases present in consensus sequence are of two types: A, G, C, T having read depth of 10 or more and lower case base a, t, g, c, n having read depth of less than 10 a/t/g/c and read depth of 0(n). Around 30 kb size consensus was generated for all samples with a range of 0.003-3.86% N’s (non-ATGC) observed in the consensus.

**Figure 2:**
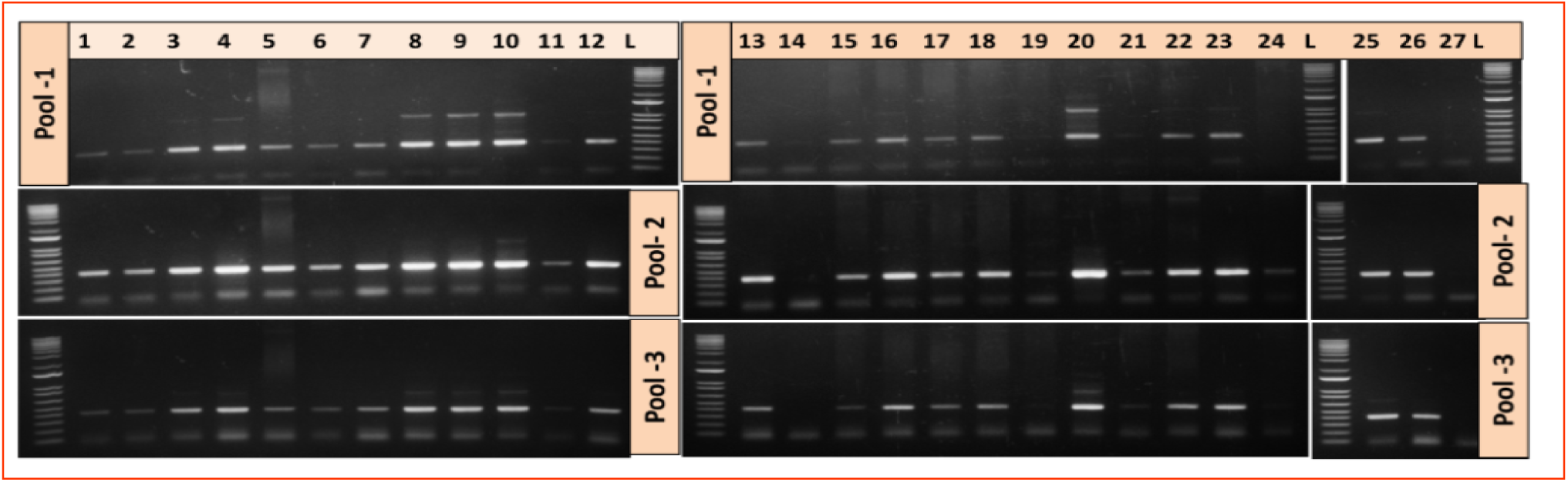
Agarose gel electrophoresis of various amplicons from primer pools targeted against SARS CoV-2, further used for nanopore sequencing.

### Molecular analysis of whole genome sequences

The SARS-CoV-2 genome sequences including viruses from different global location and within India for a period of December 2019 to May 2020, were retrieved from GISAID, EpiCov global reference web interface representative cases (Table 3). The amino acid substitutions were observed scattered throughout the genome. A total of 38 amino acid substitutions were observed compared to prototype Wuhan strain (n=38). Out of these, 24 were found in ORF1ab protein, 5 in S protein, 2 in ORF 3a, 1 each in (E, M and ORF 7a) and 4 in N protein. The unique non-synonymous variants from all samples are shown Table 4.

**Table 4:** Amino acid substitutions observed with Prototype strain isolated from Wuhan (MN908947) and COVID 19 Whole genome sequences deciphered in this study

| Amino acid Position | Gene Name | MN908947 | Virus Sequenced in this study |
| --- | --- | --- | --- |
| 27 | Orf1ab | L | F |
| 453 | Orf1ab | I | T |
| 690 | Orf1ab | A | V |
| 1158 | Orf1ab | P | S |
| 1515 | Orf1ab | S | F |
| 1534 | Orf1ab | S | I |
| 1589 | Orf1ab | T | I |
| 1597 | Orf1ab | T | I |
| 1812 | Orf1ab | A | D |
| 2016 | Orf1ab | T | K |
| 2032 | Orf1ab | D | Y |
| 2586 | Orf1ab | V | G |
| 3652 | Orf1ab | M | I |
| 3811 | Orf1ab | Y | D |
| 4207 | Orf1ab | T | I |
| 4394 | Orf1ab | A | V |
| 4489 | Orf1ab | A | V |
| 4661 | Orf1ab | D | N |
| 4696 | Orf1ab | D | G |
| 4715 | Orf1ab | L | P |
| 4721 | Orf1ab | L | I |
| 5041 | Orf1ab | S | L |
| 6083 | Orf1ab | P | S |
| 6471 | Orf1ab | Q | K |
| 583 | S | E | D |
| 614 | S | D | G |
| 884 | S | S | F |
| 929 | S | S | T |
| 943 | S | S | P |
| 81 | ORF 3a | C | R |
| 155 | ORF 3a | D | Y |
| 68 | E | S | F |
| 214 | M | S | I |
| 104 | ORF 7a | V | F |
| 13 | N | P | L |
| 18 | N | G | S |
| 139 | N | L | F |
| 378 | N | E | V |

Important non-synonymous variants includes RdRp: A97V observed in 7 strains including virus introductory index cases from three different districts, N: P13L observed in 4 strains, NSP3: T2016K observed in 7 strains including introductory index cases from three different districts. Five amino acid substitutions observed in spike protein viz., E583D in a strain from Datia, D614G observed in 17 strains from Morena, Datia, Gwalior, Ashoknagar, S884F in one strain from Morena, S929T in one strain from Ashoknagar and S943P observed in one strain of introductory index case from Gwalior district. Amino acid substitutions D614G was also observed in one of fatal case sequenced in this study.

### Phylogenetic Analysis

Whole genome based phylogenetic analysis conducted for SARS CoV-2 complete genome (29.89 Kb, n= 37) retrieved from GISAID web interface includes the representatives from all location at globe including Asia, Europe, Americas sampled between December 2019 to May 2020. The detailed molecular phylogenetic analysis revealed circulation of A1-A4 and B clades globally. Indian viruses were found to belong to multiple evolutionary genetic clades (A2a, A3, A4 and B). However, the Indian viruses sequenced in this study were found belong to three clades viz., A2a, A4 and B. Majority of virus were clustered in clade A2a (n=17), whereas seven and two viruses were found belong to clade A4 and clade B respectively. The detailed phylogeny was depicted in Fig 3.

**Figure 3:**
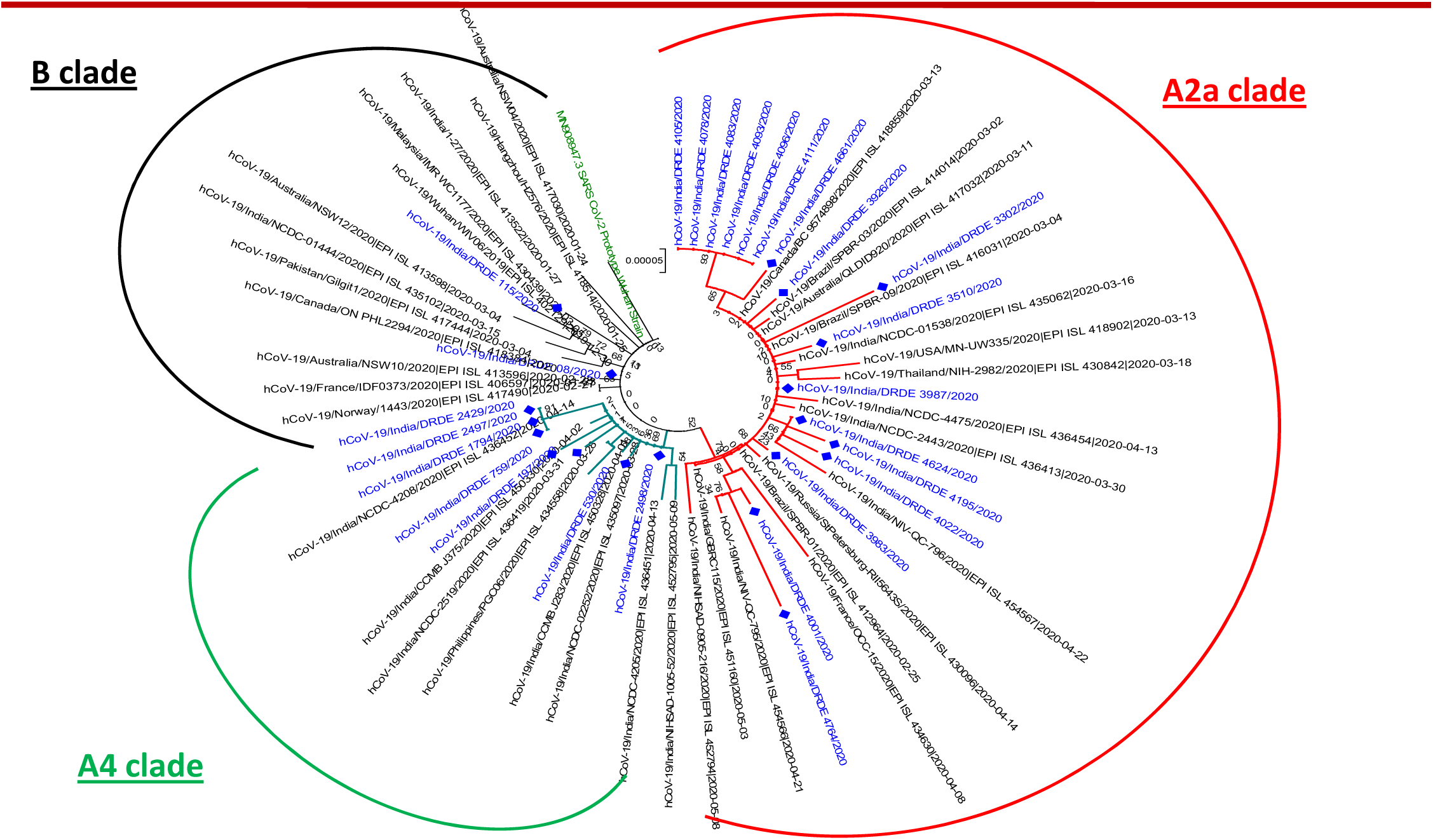
Phylogenetic analysis of Indian SARS CoV-2 sequenced in this study. The virus sequence in this study were highlighted with blue color.

## DISCUSSION

SARS-CoV-2 the etiology of COVID pandemic has been spreading throughout the world at a very rapid pace. The disease was first reported in Wuhan, China in December 2019. The rapid transmission and expansion of geographical territories has led to devastating effect on global health and economy. India is also witnessing one of the fastest growing COVID-19 infections in the world. The early lockdown in India resulted in a pause in rise of new infections. However following the mass movement of people across the nation, there is an unprecedented expansion of pandemic in different parts. The sudden emergence of SARS-CoV-2 in China is not well understood. Several reports have traced the evolutionary origins of SARSCoV-2 to SARS CoV from bats (Zhou et al, 2020) and Pangolins (Lam et al 2020). There is also a renewed interest to understand the evolutionary path being followed by SARS CoV-2 to undertake effective control and mitigation measures. In spite of large number of reported cases in India, the genomes of few Indian circulating viruses are available. Further, there is a paucity of sequence data on the genetic evolution of SARS-CoV-2 circulating in central India.

We have been investigating COVID-19 suspected cases from mid of March to May 2020, where 136 cases were found positive from central India region (10 different districts including whole Chambal region from Madhya Pradesh state). We report the sequencing of 26 whole genomes of SARS-CoV-2 viruses directly from infected patients, from ongoing COVID-19 pandemic in India.

The length of RNA genomes was found to be 29.89 kb. SARS-CoV-2, being a RNA (ribonucleic acid) virus is highly prone to mutations. A number of mutations including non synonymous substitutions (responsible for change in protein) were identified across the viruses sequenced in this study. These mutations were spread across the genome including in ORF1ab, spike, matrix, envelope and nucleoprotein regions. High evolutionary activity has been recorded in some regions (district Morena, Shivpuri, Gwalior, Datia) of the genome. An important substitution (Spike: D614G) was identified in 17 SARS-CoV-2 sequenced in this study. D614G mutation in spike protein is an interesting substitution and has been reported with increased tally [Korber et al., 2020; Chnadrika et al., 2020]. Structurally, this mutation is located in the S1 subunit that also contains the RBD domain. Although present outside the functional region, the proximity of D614G around S1 cleavage site implicates an important change in the local environment. In addition, the key mutation in spike protein (D614G) also involves loss of the charged group. These mutations that lead to positively charged groups may cause more severe structural and functional effects. Recently, this mutation has been reported to make the virus more infectious. This mutation was further linked to higher number of cases and significant mortality rate in Europe (Pachetti et al, 2020). *In vitro* studies also seem to support the hypothesis of increased transmissibility (Korber et al, 2020; Hu et al, 2020). The mutation was also recorded in one of fatal cases sequenced in this study. Presence of this mutation in majority of SARS-CoV-2 sequenced in this study indicates towards the circulation of this highly infectious virus variant in central part of India.

Of notable other significance is the P13L variant (C28311T) in the Nucleocapsid protein which is required for the viral entry into the cells (Maitra et al., 2020). The amino acid substitution was recorded in 4 viral strains, including two index cases from (Shivpuri and Morena) were recorded in samples sequenced in this study. The N protein mutations reside in the SR-rich region involved in viral capsid formation. Another important amino acid substitution P323L (RNA dependent RNA polymerase protein) was recorded in 16 viral strains sequenced in this study. This amino acid substitution is also a specific mutation for clade A2a.

The first index case (A) was identified from district Shivpuri from a person with travel history to Hyderabad in southern India. This led to emergence of the virus in this region, prior to which only 10 cases were confirmed from the whole state. During the first lockdown period another important index case (B) was reported from district Morena having travel history to Dubai. This index case led to emergence of SARS CoV-2 from district Morena. The index cases from district Sheopur and Gwalior was traced to patients having travel history to Indore and United States. Since these introductory events happened during stringent lockdown period, the infections were contained rapidly through implementation of contact tracing, quarantine and isolation.

Following the lifting of lockdown, spread of infections was reported from different districts with a remarkable positivity in whole state. The highest number of case positivity (68%) reported from district Morena in post lock down period, primarily due to reverse migration of workforce. The detailed analysis revealed multiple introduction events with diverse geographical linkage. A sudden rise in cases was observed which indicates a rapid evolution of virus in central India region. Phylogenetic analysis revealed a very distinct cladding pattern of virus emergence and evolution in different districts. The introductory index cases were found belong to clade B, whereas, virus sequence from same district were found belong to clade A4/ A2a after a gap of one month during post lock down (Unlock 1).

Majority of Indian viruses sequenced in this study were found belong to clade A2a (n=17). This is a clade having travelers importation links to Italy, UK, France. Another n=7 viruses clustered in clade A4 comprises viral entries from Southeast Asia and central Asia. The index case from district Morena, Sheopur, Shivpuri and Gwalior was having a travel history from Dubai, Indore, Hyderabad and USA. These initial index cases were found belong to separate clusters (Clade B). The further expansions in viral emergence in these districts were found with clear shift to worldwide circulating A2a and A4 clades. The independent grouping with different clusters of infection with region was also observed.

In summary, this is the first detailed genomic signature analysis of SARS-CoV-2 emergence and expansion in the Chambal region of state Madhya Pradesh, central India. This analysis also explained the initial introductory link to further virus evolution in different districts in pre and post lockdown period. Continuous monitoring of genomic architecture of SARS-CoV-2 for possible mutations is crucial for success of diagnostics, vaccine and therapeutics. This will be useful in mapping the spread and evolution of COVID-19 in future and can aid in microbial forensics investigation.

## ACKNOWLEDGEMENTS

The authors do acknowledge GISAID for sharing the genomic sequences in public domain and other contributors for SARS-CoV-2 genomic data. The authors would like to thank the financial aid provided by Defence Research & Development Organization (DRDO), Ministry of Defence, Government of India for this work. We would also thanks the chief medical health officer of district Morena, Gwalior, Shivpuri, Sheopur, Ashok Nagar, Guna, Datia, Bhind and Joint Director, health, Gwalior for referring samples for lab diagnosis. This manuscript is assigned DRDE accession no. DRDE/VIRO/15/2020.

## AUTHOR CONTRIBUTIONS

Shashi Sharma: RT-PCR, Sequencing, Results analysis, wrote MS,; Paban Kumar Dash: Design, Results analysis, supervision, Review; Sushil K Sharma: Sample Processing; Ambuj Srivastava: Sample Processing; Jyoti S Kumar: RT-PCR; B.S. Karothia: Sample Processing; K T Chelvam: Sample Processing; Sandip Singh: Sample Processing; Abhay Gupta: Sample Processing; Ram Govind Yadav: Sample Processing; Ruchi Yadav: Sample Processing; Greshma TS: Sample Processing; Pramod Kushwah: Sample Processing; Ravi Bhushan: Sample Processing; D.P. Nagar: Sample Processing; Manvendra Nandan: Sample Processing; Subodh Kumar: Supervision, Review; Duraipandian Thavaselvam: Supervision, Review; Devendra Kumar Dubey:Supervision, Review. All the authors read and approved the final manuscript.

## DECLARATION OF INTERESTS

Authors declare no conflict of interests with the present study.

## Notes

### Competing Interest Statement

The authors have declared no competing interest.

